# The uniqueness of the Brazilian Yellow Fever Virus. Are the vaccines less effective against it?

**DOI:** 10.1101/2023.05.05.539667

**Authors:** Prabhudutta Mamidi, Mahendra Gaur, Baijayantimala Mishra, Bharat Bhusan Subudhi, Sutapa Rath, Monalisa Mohanty

## Abstract

The years 2016-2019 witnessed the expansion of Yellow Fever Virus (YFV) circulation area in Brazil. Deforestation and mutations in the nsp genes were presumed responsible for the reemergence of YFV in brazil. However, our data analysis involving 200 clinical isolates worldwide including vaccine strains showed two amino acids substitutions exclusively in the Brazilian strains at critical positions (V318A and I335M) in the ED III domain of E protein. Our molecular dynamics analysis involving reference and mutant structures suggests that the mutations significantly changed the conformational rearrangement and folding of the YFV-EDIII domain as revealed by RMSF, PCA, Porcupine plots and secondary structure analysis. Briefly, the regions 324-330, 339-346 and 347-359 had a higher fluctuation than the wild, while regions 313-318 and 379-384 had a lower fluctuation. As reported earlier that the residues E325 and E380 are part of vaccine epitopes and any mutations in these residues reduced the binding of vaccine generated mabs to the epitopes. Hence, it may be speculated from our porcupine plot data analysis that that the commercial YFV vaccines may not be fully effective against the Brazilian strains in particular owing to the fluctuations of these residues. However, future investigation is required to validate the observations.

## Yellow Fever Virus :-a brief introduction

Yellow Fever Virus (YFV) is an insect-borne virus belonging to the *flaviviridae* family. It is spread due to *Aedes* or *Hemagogus* mosquitoes. It has two cycles, sylvatic and urban. Yellow Fever (YF) can be asymptomatic or symptomatic with fever, myalgia, back pain and prostration and in some cases lead to multi-organ failure dominated by hepato-renal failure and profound jaundice. YF still remains a public health threat in the human populations of Africa and South America with a high Case Fatality Rate (CFR) ranging from 40% - 60% especially in South America.^1^ According to the Pan American Health Organization (PAHO), the South American countries that reported the highest numbers of cases of YF during 1960–2019 were Brazil (3829 cases), Peru (3189 cases), Bolivia (1546 cases) and Colombia (701 cases).^2^

## Re-emergence of YFV in Brazil: possible reasons

It is assumed that the YFV and its vector *Aedes aegyptii* was introduced in brazil through the slave trading ships from West Africa during the early colonization period. In Brazil, the first YF epidemic was recorded in 1685. However, the last two decades have witnessed the expansion of YFV circulation area in the country. The extensive re-emergence of YF in Brazil started in late 2016, and, according to data from the Ministry of Health, 2237 human cases of YF and 759 deaths were recorded between December 2016 and June 2019.^2^ From the above data, one needs to speculate on the reasons of this re-emergence:

Firstly, the recent re-emergence of YFV showed that the majority of the population affected by YF (82.8% during 2017–2018) were males of economically active age range and were residents of rural areas in proximity to forest sites.^2-3^ Thus, it can be speculated that deforestation may be a factor that enhanced human exposure to fragmented forest areas thereby increasing the risk of YFV spread to urban environments through YFV sylvatic cycles. Moreover, experimental studies also suggest that YFV has the potential to adapt to *A. albopictus*, which is an opportunistic species that circulates between urban and peri-urban habitats. This may also be one of the reasons for the re-emergence.^4^

Secondly, some studies performed in Brazil(State of Minas Gerais) have highlighted the absence of neutralizing antibodies against YFV in individuals from rural and urban areas. This was also considered as a reason for the recent epidemic in the region. Furthermore, some individuals from the same area with proven vaccination record against YFV had no neutralizing antibodies. The above statement is interesting however, in one of the studies conducted in 2019, it has been observed that that the vaccinees, who had seroconverted following primary vaccination (17DD-YF vaccine) with sub-doses (10447IU;3013IU;587IU respectively) in 2009, presented similar neutralizing antibodies levels 8 years after primary vaccination as compared to the reference full dose (27476IU).^5^ This shows that there might be some unexplored reasons behind the absence of neutralizing antibodies in the individuals during the Brazilian epidemic.

## Are the vaccines less effective against the Brazilian YFV strain ?

The live attenuated 17D vaccine against YFVwas developed in the 1930s and currently, there are three 17D substrains in production; 17DD manufactured in Brazil, 17D-213 manufactured in Russia, and 17D-204 manufactured in China, France, Senegal, and the USA. Four of the vaccines (Brazil, France, Russia, and Senegal) are prequalified and used internationally for WHO/UNICEF vaccination campaigns.^6^

However, in order to answer the above question, we had to explore the YFV E protein sequences. Now, in case of flaviviruses, entry into the target cell is mainly dependent on the E protein interaction with its cognate receptor. The E protein of YFV consists of three ecto-domains (E-DI, E-DII, and E-DIII).Out of these, EDIII contains important linear antigenic epitopes that directly interact with potent neutralizing antibodies.^7^ First, we aligned the YFV ED III protein sequences derived from patient samples worldwide from NCBI. It was observed that there were two amino acid substitutions in the most conserved position that were unique to the brazilian strains namely V318A and I335M and these were absent in rest of the sequences worldwide as shown in Supp Fig S1. This prompted us to explore the effect of these substitutions on ED III structure. Therefore, The crystal structure (PDB: 2JQM; From Met287 to Lys398) of yellow fever virus (YFV) E protein’s domain III (YFV-EDIII) was retrieved from Protein data bank (PDB) (https://www.rcsb.org/) and used as a reference structure,^8^ for designing the mutant structure with four substitutions (V318A, K331R, I335M and I344V) which are unique to the Brazilian strains. The K331R and I344V mutations were absent in the PDB reference structure so it was included for constructing the mutant ED III structure.

## How is the mutant structure generated using homology modeling?

### 1. Structure preparation

During the structure preparation, the missing hydrogen atoms were added, missing side chains and loops were filled using the Prime and protonation states at pH 7.0 was generated using the PROPKA of the Schrödinger platform. For generating mutant variant, the residuesVal318, Lys331, Ile335 and Ile344 was mutated to Ala318, Arg331, Met335 and Val344 respectively. The total number of atoms in both the wild and mutant was 1688.

### 2. Molecular Dynamics simulation and post-dynamic analysis

To evaluate the effect of the point mutations on the E protein domain’s structural stability, we employed the 100 ns scale molecular dynamics (MD) simulations of the domain III in the water environment at 300 K using the GROMACS suite v2021.1 with charmm36-jul2021 force field.^9-10^ Both the variants (wild and mutant) of E protein domain III (EDIII) were solvated in a cubic periodic box with Tip3P charm-modified water model. Systems were neutralized by adding NaCl salt of concentration 0.15 M followed by energy minimization of a maximum of 50,000 steps using the steepest descent method. The systems were then subjected to NVT (constant number of particles, volume and temperature) and NPT (constant number of particles, pressure and temperature) equilibrations at pressure and temperate of 1 bar and 300 K respectively. To constraint all the bonds and angles, LINear Constraint Solver (LINCS) algorithm was used,^11^ whereas the Verlet algorithm was used to integrate Newton’s equations of motion. The temperature and pressure coupling in each equilibrations steps was controlled by V-rescale (modified Berendsen thermostat)and Parrinello-Rahman method respectively. However, in NVT equilibrations no controller was used for pressure coupling. The time evolution of trajectories was recorded every 10 ps for 100 ns. After completion of 100 ns scale simulation, the trajectories were comparatively analyzed for root mean square deviation (RMSD), root mean square fluctuation (RMSF), solvent accessible surface area (SASA) and Number of intramolecular hydrogen bonds formations using the in-built module of the GROMACS.

### 3. Clustering and structure alignment

The generated molecular dynamics simulation trajectories were clustered using a Python package TTClust to extract the representative frame (medoids) for each cluster.^12^ The TTClust uses elbow statistics to find the optimal number of clusters based on RMSD-based distance between frames. The medoids of the most populated cluster from wild and mutant clustering were then superimposed to observe the RMSD difference between them.

### 4. Essential Dynamics analysis

To reduce the complexity of the trajectories data and extract the biologically significant motion in wild and mutant YFV-EDIII structure over the course of 100 ns simulation, we performed the Principal component analysis (PCA) on Cα atoms using the MODE-TASK.^13^ Further, the movement and amplitude of the motion in Cα atoms throughout the simulation in the wild and mutant structure, the porcupine plots of principal component 1 and 2 data were generated using the ProDy plugin (https://github.com/prody/ProDy) of VMD molecular visualization program.^14^

### 5. Secondary structure elements (SSE) analysis

To observe the changes in the secondary structure of each residue for all frames in wild and mutant structure throughout the simulation, we used the VMD plugin ‘Timeline’ to calculate the secondary structure of each frame. The extracted data were then plotted by ggplot2 R package.^15^

What does the data reveal? Are there any significant changes in wild type and mutant structure?

## Post-dynamic analysis of wild and mutant YFV-EDIII

The wild and mutant structure of YFV-EDIII was subjected to 100 ns MD simulation to observe the possible impact of four substitutions (V318A, K331R, I335M and I344V) on structural and conformational changes of the domain. The RMSD analysis of Cα atoms for both wild and mutant variants, suggests an initial escalation by 4.3 Å upto 30 ns and then decrease. After 30 ns, it showed stability (RMSD of 2.0-3.0 Å), indicating convergence of both structures.(Figure 1A). According to RMSF analysis, in both variants of YFV-EDIII domain, almost all residues significantly contributed to its fluctuation. The region 324-330, 339-346 and 347-359 showed a higher fluctuation than the wild, suggesting an increase in the flexibility of these regions compared to wild structure. Whereas, regions 313-318 and 379-384 showed a lower fluctuation, implying lesser flexibility and more compactness (Figure 1B). These fluctuations may affect the binding with the host target proteins. Solvent-accessible surface area (SASA) was analyzed to observe the changes in solvent-accessible bimolecular surface area (solvent accessibility) of the mutant structure compared to wild structure. The mutant structure showed a comparatively lower average SASA value (60.5±1.4 nm^2^)than the wild structure (64.08±1.4 nm^2^) in the stable region after 30 ns(Figure 1C).The lower value of SASA suggests the hydrophobic compactness and increased stability of the structure. The structural changes over the simulation time could lead to the rearrangementof intramolecular hydrogen bond formations. The average number of hydrogen-bond formation in the wild and mutant were 51.07±3.67 and 53.17±4.81 respectively (Figure 1D). The more number of H-Bonds in the mutant structure suggest the stability of the protein over the simulation time.

**Figure 1:**
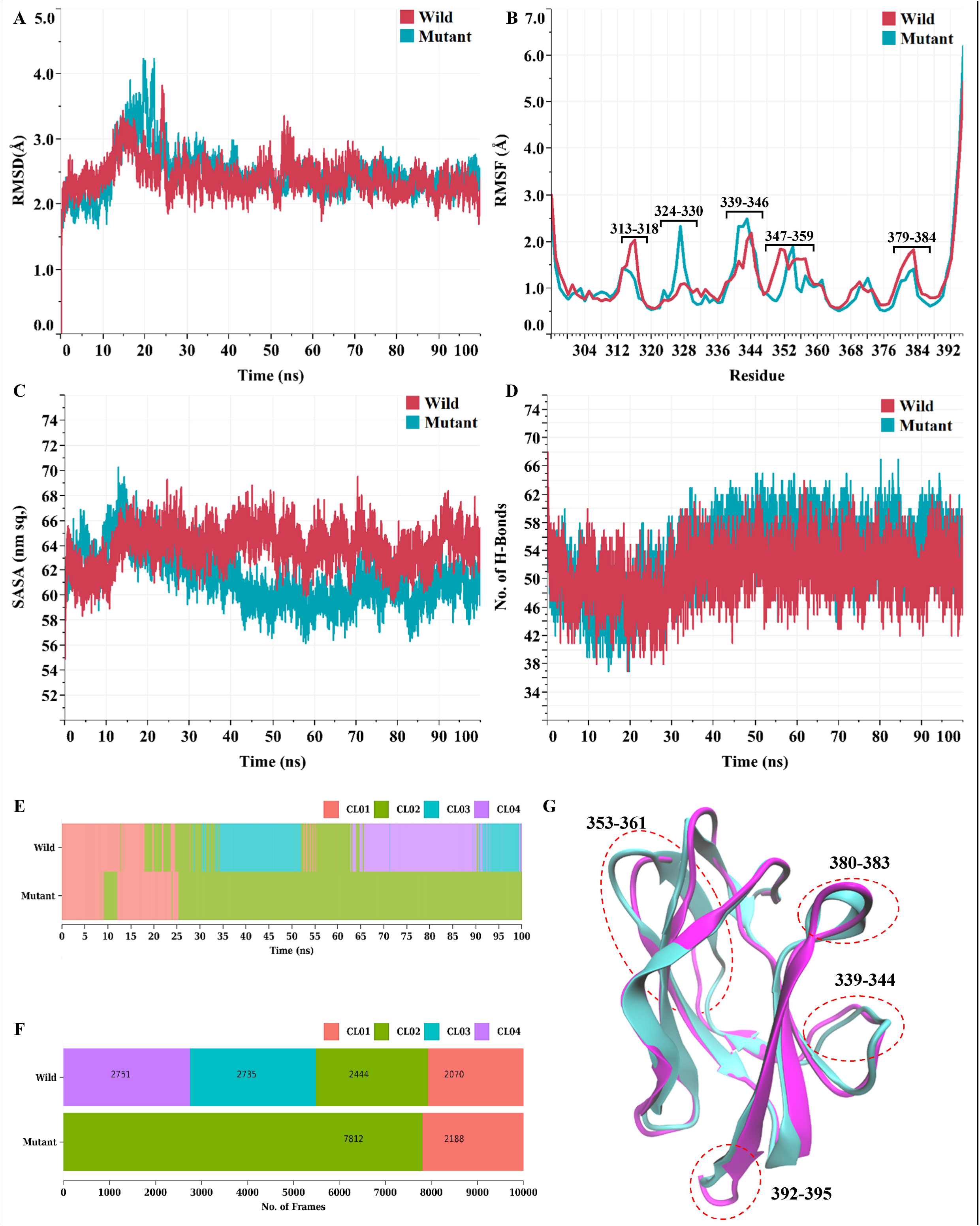
Molecular dynamics simulation trajectories analysis, clustering and 3D superimposition of wild and mutant E protein domain III (EDIII; Residue 296 to 395) variants over 100 ns simulation time. (A) Root Mean Square Deviation (RMSD) value of Cα atoms. (C) Solvent Accessible Surface Area (SASA) of the whole domain. (D) The number of intramolecular hydrogen bond formations. (E) 2D linear projection of clusters over 100 ns simulation. Each bar line represents a frame. (F) Staked bar plot depicting the overview of the cluster size. The size of the population for each cluster is represented as a label on each staked bar. (G) 2D superimposition of medoids structure extracted from the most populated cluster. The structure of wild (in magenta color) and mutant (in cyan color) is represented in cartoon form. Thirty-four residues (296-298, 303, 313, 323, 326, 327, 331, 339-344, 353-361, 371, 380-383, 388, 392-395) show the RMSD ≥2.0 Å as compared to the wild structure. A few of the regions are highlighted in the dashed red colored circle.

## Conformational heterogeneity between wild and mutant

The TTClust was used for clustering the ensembles of both wild and mutant conformations samples during 100 ns simulations. The clustering segregated the ensembles of both wild and mutant trajectories into four and two optimal clusters for wild and mutant respectively by elbow statistics based on Cα RMSD(Figure 1E and F). The medoids of the most populated cluster from wild and mutant clustering were superimposed to observe the 3D structural deviation. An RMSD value of 4.9 Å for mutant structure indicates the larger changes in the intra-atomic bond network and secondary structure of residues (Figure 1G). Further, thirty-four residues (296-298, 303, 313, 323, 326, 327, 331, 339-344, 353-361, 371, 380-383, 388, 392-395) showed RMSD ≥2.0 Å as compared to the wild structure (Supp. Figure S2).

## Essential dynamics analysis reveals the significant changes in the motion of Cα atoms due to mutations

To study the impacts of mutations on the changes in the dynamics of motions of Cα atoms, we also performed principal component analysis using cartesian coordinates and singular value decomposition (SVD) approach. From the analysis of the collective movement of Cα atoms,the scatter plot generated for the wild (Fig. 2A) and mutant (Figure 2B)indicated a significant difference between the wild and mutant. The trace values observed for the wild and mutant were 1.87 nm^2^ and 1.85 nm^2^ respectively, which suggest less 3D structural conformations for the mutant than the wild (Figure 2C). In the mutant, the first three components explained the 57.54% variance as compared to the mutant which was 54.63% (Figure 2D).Further, to study the quantitative motion in Cα atoms throughout the simulation captured by first two principal components, porcupine plots was generated using the ProDy plugin of VMD program. The directions and length of the arrow represent the directions and amplitude of the motion. The motion of region 341-345 in the mutant domain shows higher motion on both components (PC1 and PC2) as compared to the wild domain (Figure 2E and F). From porcupine plots, the changes in directions and amplitude of the motions reveal the conformational changes in the mutant as compared to the wild structure over the 100 ns simulations. The detailed fluctuations in Cα atoms of each residue is represented in supplementary Table S1.

**Figure 2:**
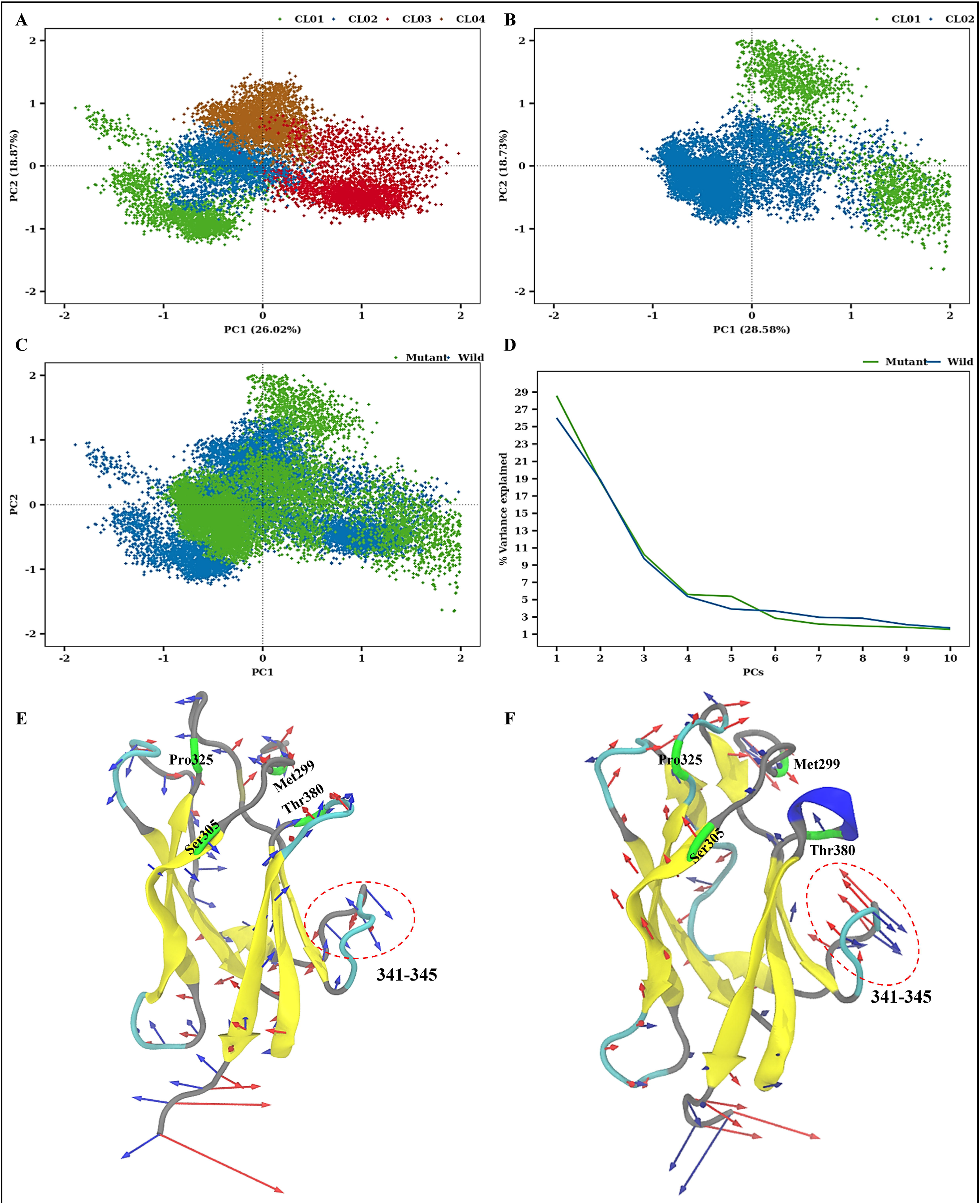
Combined essential dynamics (ED) analysis over 100 ns simulation time which separates the motions of the of wild and mutant E protein domain III (EDIII; Residue 296 to 395) variants into principal modes (PC1 vs PC2). (A) 2D projections of wild EDIII trajectory in essential subspace. (B) 2D projections of mutant EDIII trajectory in essential subspace. (C) Combined 2D projections of wild and mutant EDIII trajectory in essential subspace. (D) Percentage of variance of top ten principal components/modes for wild and mutant EDIII trajectories. Porcupine plots of significant motion obtained from PCA across the PC1 (Blue) and PC2 (Red) in wild (E) and mutant (F) trajectories. The four residues of EDIII epitopes that was identified using monoclonal antibodies (mAbs) were highlighted in green colour and labelled accordingly. The fluctuations in Cα atoms above 1.0 Å in in the form red (PC1) and blue (PC2) coloured cones is drawn. The length of the cone is scaled by 2.0 Å RMSD and depicts the direction as well the amplitude/magnitude of the motion. The structure is represented in cartoon form where α-helix by purple, 3_10_ helix by blue, π-helix by red, Extended β-sheets by yellow, β-bridge by tan, turn by cyan and coils by grey colour are shown.

## Changes in Secondary structure elements

The time evolution of the secondary structure of each residue in both wild and mutant was computed using VMD timeline plugins and plotted using the ggplot2 R package (Supp. Figure S3A and B). From the secondary structure analysis, we observed the stability of the secondary structure of each residue throughout the simulations. The region 380-384 of mutant structure changed from turn (yellow) to 3_10_-helix (red) in the last 10 ns of the simulations (Supp. Figure S3B).

Overall, our molecular dynamics analysis suggests that the four mutations significantly changed the conformational rearrangement and folding of the YFV-EDIII domain as revealed by RMSF, PCA, Porcupine plots and secondary structure analysis.

## What are the possible implications of the findings ?

Out of the five identified regions (324-330, 339-346, 347-359, 313-318 and 379-384) in this study, two have been characterized in YFV for carrying important residues for antigenicity/ neutralizing epitopes (324-330 and 379-384). In one of the recent articles published in 2022, it has been observed that the vaccine (17D-204, 17DD and 17D-213) specific epitope residues recognized by mab 411 (E-299, E305, E325 and E380) are physically located on ED III and any changes to these residues affected the binding of mabs to the epitopes.^16^ From Figure 2E and F, it seems there is a shift in rotation in residues Pro325 and Thr380 between wild type and mutant structure which may affect the binding of pre-existing mabs to these residues.

Additionaly, the region (339-359) showing the maximum fluctuation should also be considered and characterized in future to understand the complete role of the unique substitutions that are confined to the Brazilian strains only. Because ths region contains residues which are part of the low pH trimer interface domain which is important for viral entry into the host cell. However, further experimental validation is required to explore the significance of these fluctuations. Thus, it seems that the currently used commercially available YFV vaccine (17DD) in brazil may not be as effective in neutralizing YFV as expected. If this viewpoint is correct and problems are not sorted out early then brazil should be prepared for future YFV epidemics.

## Authors Contributions

PM is involved in conceptualisation, data curation, formal analysis and writing-original draft.MG is involved in data curation, formal analysis and writing original draft. BM and BBS are involved in review and editing of the draft. SR and MM are involved in data curation and editing of the draft.

## Declaration of Interests

The authors declare no conflict of Interest

## Acknowlegements

MG is supported by a fellowship funded from by Department of Biotechnology (DBT), Ministry of Science and Technology, New Delhi, India (Grant Id: BT/INF/22/SP45078/2022). The authors would like to acknowledge the Indian Council of Medical Research (ICMR), New Delhi, India (Grant ID: AMR/DHR/GIA/4/ECD-II-2020) for providing high-performance computational resources for this study.

## Notes

### Competing Interest Statement

The authors have declared no competing interest.

