## Supplemental file for "The uniqueness of the Brazilian Yellow Fever Virus. Are the vaccines less effective against it?"

**Supp Figure S1: Screen shots showing the alignment of YFV strains including the vaccine strains from all over the world**. Around 200 strains from all over the world including the four vaccine strains(17D, 17DD,17D-204, 17D-213) were aligned using the Clustal W tool. The amino acid positions unique to the brazilian strains are flagged in red at the top. The 2JQM (PDB ID) fasta sequence is taken as the reference sequence for comparison.

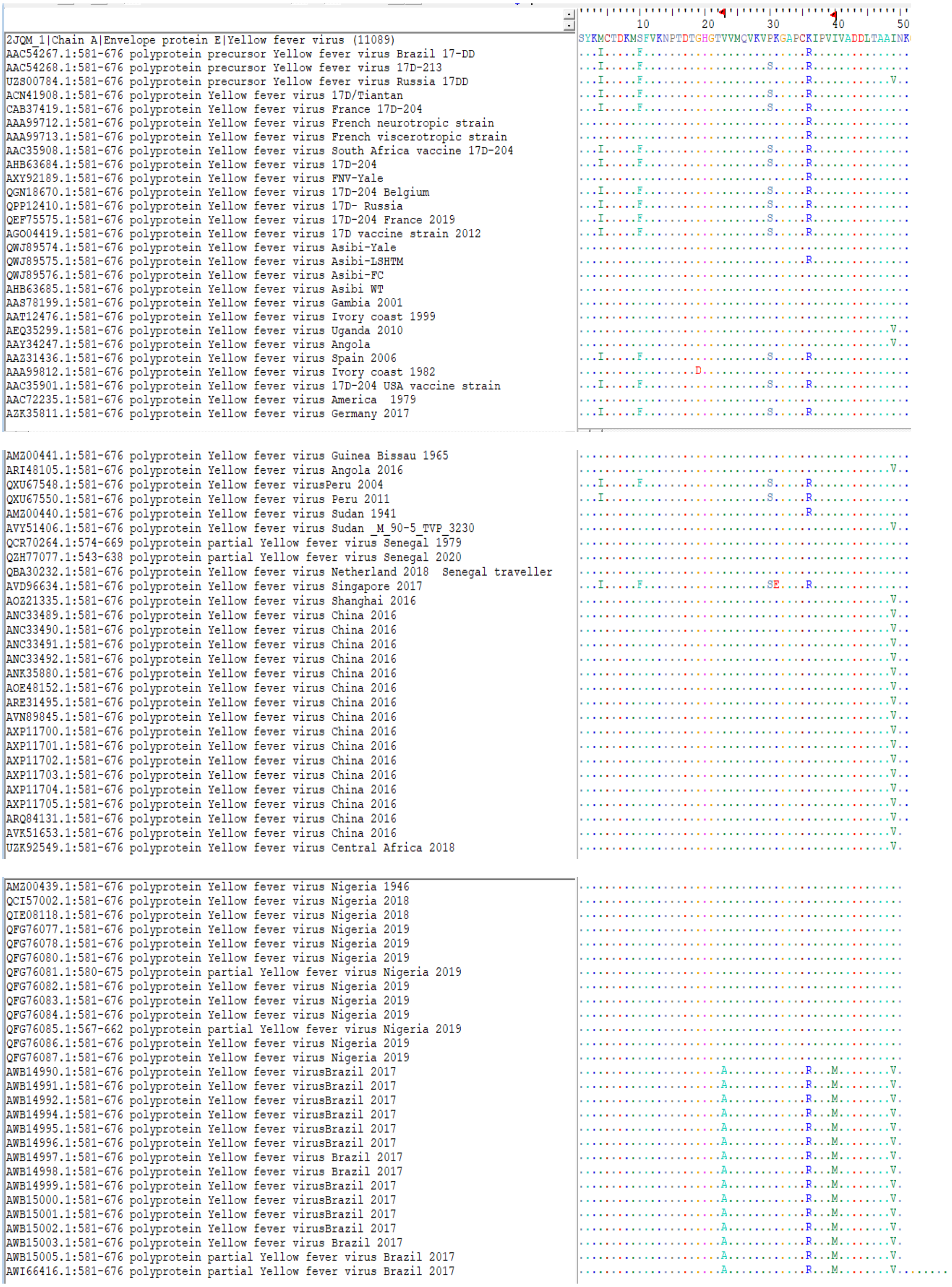

| | |
|---|---|
| ATN45408.1:581-676 polyprotein Yellow fever virus Brazil 2017 | ......................A............R...M........V........ |
| QBB02272.1:573-668 polyprotein partial Yellow fever virus Brazil 2017 | ......................A............R...M........V........ |
| AVQ67777.1:581-676 polyprotein Yellow fever virus Brazil 2017 | ......................A............R...M........V........ |
| AVQ94369.1:581-676 polyprotein Yellow fever virus Brazil 2017 | ......................A............R...M........V........ |
| AVQ94370.1:581-676 polyprotein Yellow fever virus Brazil 2017 | ......................A............R...M........V........ |
| AVQ94371.1:581-676 polyprotein Yellow fever virus Brazil 2017 | ......................A............R...M........V........ |
| AVQ94374.2:581-676 polyprotein partial Yellow fever virus Brazil 2017 | ......................A............R...M........V........ |
| QEZ88133.1:573-668 polyprotein partial Yellow fever virus Brazil 2017 | ......................A............R...M........V........ |
| QEZ88136.1:573-668 polyprotein partial Yellow fever virus Brazil 2017 | ......................A............R...M........V........ |
| QEZ88139.1:573-668 polyprotein partial Yellow fever virus Brazil 2017 | ......................A............R...M........V........ |
| QEZ88140.1:437-532 polyprotein partial Yellow fever virus Brazil 2017 | ......................A............R...M........V........ |
| QEZ88143.1:573-668 polyprotein partial Yellow fever virus Brazil 2017 | ......................A............R...M........V........ |
| QEZ88145.1:573-668 polyprotein partial Yellow fever virus Brazil 2017 | ......................A............R...M........V........ |
| ASY08185.1:581-676 polyprotein Yellow fever virus Brazil 2017 | ......................A............R...M........V........ |
| QMT62826.1:573-668 polyprotein partial Yellow fever virus Brazil 2017 | ......................A............R...M........V........ |
| ASY08187.1:581-676 polyprotein Yellow fever virus Brazil 2017 | ......................A............R...M........V........ |
| AXG65492.1:581-676 polyprotein Yellow fever virus Brazil 2017 | ......................A............R...M........V........ |
| AZA36620.1:573-668 polyprotein partial Yellow fever virus Brazil 2017 | ......................A............R...M........V........ |
| QDA77104.1:581-676 polyprotein Yellow fever virus Brazil 2017 | ......................A............R...M........V........ |
| AWF71047.1:580-675 polyprotein partial Yellow fever virus Brazil 2017 | ......................A............R...M........V........ |
| AWF71049.1:580-675 polyprotein partial Yellow fever virus Brazil 2017 | ......................A............R...M........V........ |
| AWF71051.1:580-675 polyprotein partial Yellow fever virus Brazil 2017 | ......................A............R...M........V........ |
| AWF71052.1:580-675 polyprotein partial Yellow fever virus Brazil 2017 | ......................A............R...M........V........ |
| AWF71054.1:580-675 polyprotein partial Yellow fever virus Brazil 2017 | ......................A............R...M........V........ |
| AWF71055.1:580-675 polyprotein partial Yellow fever virus brazil 2017 | ......................A............R...M........V........ |
| AWF71056.1:580-675 polyprotein partial Yellow fever virus Brazil 2017 | ......................A............R...M........V........ |
| AWF71057.1:580-675 polyprotein partial Yellow fever virus Brazil 2017 | ......................A............R...M........V........ |
| AWF71058.1:580-675 polyprotein partial Yellow fever virus Brazil 2017 | ......................A............R...M........V........ |

| | |
|---|---|
| AWF71063.1:580-675 polyprotein partial Yellow fever virus Brazil 2017 | ......................A............R...M........V........ |
| AWF71065.1:580-675 polyprotein partial Yellow fever virus Brazil 2017 | ......................A............R...M........V........ |
| AWF71066.1:580-675 polyprotein partial Yellow fever virus Brazil 2017 | ......................A............R...M........V........ |
| AWF71067.1:580-675 polyprotein partial Yellow fever virus Brazil 2017 | ......................A............R...M........V........ |
| AWF71068.1:580-675 polyprotein partial Yellow fever virus Brazil 2017 | ......................A............R...M........V........ |
| AWF71069.1:580-675 polyprotein partial Yellow fever virus Brazil 2017 | ......................A............R...M........V........ |
| AWF71070.1:580-675 polyprotein partial Yellow fever virus Brazil 2017 | ......................A............R...M........V........ |
| AWF71071.1:580-675 polyprotein partial Yellow fever virus Brazil 2017 | ......................A............R...M........V........ |
| AWF71072.1:580-675 polyprotein partial Yellow fever virus Brazil 2017 | ......................A............R...M........V........ |
| AWF71078.1:573-668 polyprotein partial Yellow fever virus Brazil 2003 | ......................A............R...M........V........ |
| AWF71079.1:573-668 polyprotein partial Yellow fever virus Brazil 2003 | ......................A............R...M........V........ |
| AWF71081.1:573-668 polyprotein partial Yellow fever virus Brazil 2017 | ......................A............R...M........V........ |
| AWF71082.1:573-668 polyprotein partial Yellow fever virus Brazil 2017 | ......................A............R...M........V........ |
| AWF71083.1:573-668 polyprotein partial Yellow fever virus Brazil 2017 | ......................A............R...M........V........ |
| AWF71085.1:581-676 polyprotein partial Yellow fever virus Brazil 2017 | ......................A............R...M........V........ |
| AWF71089.1:573-668 polyprotein partial Yellow fever virus Brazil 2017 | ......................A............R...M........V........ |
| AWF71091.1:573-668 polyprotein partial Yellow fever virus Brazil 2017 | ......................A............R...M........V........ |
| AXN92351.1:573-668 polyprotein partial Yellow fever virus Brazil 2017 | ......................A............R...M........V........ |
| AZA36619.1:573-668 polyprotein partial Yellow fever virus Brazil 2017 | ......................A............R...M........V........ |
| AZA36622.1:573-668 polyprotein partial Yellow fever virus Brazil 2017 | ......................A............R...M........V........ |
| UVG67830.1:573-668 polyprotein partial Yellow fever virus Brazil 2018 | ......................A............R...M........V........ |
| UVG67831.1:573-668 polyprotein partial Yellow fever virus Brazil 2018 | ......................A............R...M........V........ |
| UVG67832.1:573-668 polyprotein partial Yellow fever virus Brazil 2018 | ......................A............R...M........V........ |
| UVG67833.1:573-668 polyprotein partial Yellow fever virusBrazil 2018 | ......................A............R...M........V........ |
| UVG67834.1:573-668 polyprotein partial Yellow fever virus Brazil 2018 | ......................A............R...M........V........ |
| UVG67835.1:573-668 polyprotein partial Yellow fever virus Brazil 2018 | ......................A............R...M........V........ |
| UVG67836.1:573-668 polyprotein partial Yellow fever virus Brazil 2018 | ......................A............R...M........V........ |
| UVG67837.1:573-668 polyprotein partial Yellow fever virus Brazil 2018 | ......................A............R...M........V........ |

| | |
|---|---|
| UVG67838.1:573-668 polyprotein partial Yellow fever virus Brazil 2018 | ......................A............R...M........V........ |
| UVG67840.1:573-668 polyprotein partial Yellow fever virus Brazil 2018 | ......................A............R...M........V........ |
| QEZ88135.1:573-668 polyprotein partial Yellow fever virus Brazil 2018 | ......................A............R...M........V........ |
| QEZ88150.1:573-668 polyprotein partial Yellow fever virus Brazil 2018 | ......................A............R...M........V........ |
| QEZ88151.1:573-668 polyprotein partial Yellow fever virus Brazil 2018 | ......................A............R...M........V........ |
| QEZ88153.1:573-668 polyprotein partial Yellow fever virus Brazil 2018 | ......................A............R...M........V........ |
| QHB50136.1:581-676 polyprotein Yellow fever virus Brazil 2018 | ......................A............R...M........V........ |
| QHB50137.1:581-676 polyprotein Yellow fever virus Brazil 2018 | ......................A............R...M........V........ |
| QHB50138.1:581-676 polyprotein Yellow fever virus Brazil 2018 | ......................A............R...M........V........ |
| QHB50139.1:581-676 polyprotein Yellow fever virus Brazil 2018 | ......................A............R...M........V........ |
| QHB50140.1:581-676 polyprotein Yellow fever virus Brazil 2018 | ......................A............R...M........V........ |
| QHB50141.1:581-676 polyprotein Yellow fever virus Brazil 2018 | ......................A............R...M........V........ |
| QHB50142.1:581-676 polyprotein Yellow fever virus Brazil 2018 | ......................A............R...M........V........ |
| QHB50143.1:581-676 polyprotein Yellow fever virus Brazil 2018 | ......................A............R...M........V........ |
| QHB50144.1:581-676 polyprotein Yellow fever virus Brazil 2018 | ......................A............R...M........V........ |
| QHB50145.1:581-676 polyprotein Yellow fever virus Brazil 2018 | ......................A............R...M........V........ |
| QHB50146.1:581-676 polyprotein Yellow fever virus Brazil 2018 | ......................A............R...M........V........ |
| QHB50147.1:581-676 polyprotein Yellow fever virus Brazil 2018 | ......................A............R...M........V........ |
| QHB50148.1:581-676 polyprotein Yellow fever virus Brazil 2018 | ......................A............R...M........V........ |
| QHB50149.1:581-676 polyprotein Yellow fever virus Brazil 2018 | ......................A............R...M........V........ |
| QHB50150.1:581-676 polyprotein Yellow fever virus Brazil 2018 | ......................A............R...M........V........ |
| QHB50151.1:581-676 polyprotein Yellow fever virus Brazil 2018 | ......................A............R...M........V........ |
| QHB50152.1:581-676 polyprotein Yellow fever virus Brazil 2018 | ......................A............R...M........V........ |
| QHB50153.1:581-676 polyprotein Yellow fever virus Brazil 2018 | ......................A............R...M........V........ |
| QHB50154.1:581-676 polyprotein Yellow fever virus Brazil 2018 | ......................A............R...M........V........ |
| QHB50155.1:581-676 polyprotein Yellow fever virus Brazil 2018 | ......................A............R...M........V........ |
| QHB50156.1:581-676 polyprotein Yellow fever virus Brazil 2018 | ......................A............R...M........V........ |
| QHB50157.1:581-676 polyprotein Yellow fever virusBrazil 2018 | ......................A............R...M........V........ |

| | |
|---|---|
| QHB50158.1:581-676 polyprotein Yellow fever virus Brazil 2018 | ......................A............R...M........V........ |
| QHB50159.1:581-676 polyprotein Yellow fever virus Brazil 2018 | ......................A............R...M........V........ |
| QHB50160.1:581-676 polyprotein Yellow fever virus Brazil 2018 | ......................A............R...M........V........ |
| QHB50161.1:581-676 polyprotein Yellow fever virus Brazil 2018 | ......................A............R...M........V........ |
| QHB50162.1:581-676 polyprotein Yellow fever virus Brazil 2018 | ......................A............R...M........V........ |
| QHB50163.1:581-676 polyprotein Yellow fever virus Brazil 2018 | ......................A............R...M........V........ |
| QHB50164.1:581-676 polyprotein Yellow fever virus Brazil 2018 | ......................A............R...M........V........ |
| QHB50165.1:581-676 polyprotein Yellow fever virus Brazil 2018 | ......................A............R...M........V........ |
| QHB50166.1:581-676 polyprotein Yellow fever virus Brazil 2018 | ......................A............R...M........V........ |
| QHB50167.1:581-676 polyprotein Yellow fever virus Brazil 2018 | ......................A............R...M........V........ |
| QHB50168.1:581-676 polyprotein Yellow fever virus Brazil 2018 | ......................A............R...M........V........ |
| QHB50169.1:581-676 polyprotein Yellow fever virus Brazil 2018 | ......................A............R...M........V........ |
| QHB50170.1:581-676 polyprotein Yellow fever virus Brazil 2018 | ......................A............R...M........V........ |
| QHB50171.1:581-676 polyprotein Yellow fever virus Brazil 2018 | ......................A............R...M........V........ |
| QBK94341.1:581-676 polyprotein Yellow fever virus Brazil 2018 | ......................A............R...M........V........ |
| QBK94342.1:581-676 polyprotein Yellow fever virus Brazil 2018 | ......................A............R...M........V........ |
| UO885857.1:581-676 polyprotein Yellow fever virus Brazil 2018 | ......................A............R...M........V........ |
| QWJ73458.1:581-676 polyprotein Yellow fever virus Brazil 2018 | ......................A............R...M........V........ |
| QWJ73459.1:581-676 polyprotein Yellow fever virus Brazil 2018 | ......................A............R...M........V........ |
| AFH35034.1:581-676 polyprotein Yellow fever virus Brazil 1981 | ......................A............R...M........V........ |
| AFH35037.1:581-676 polyprotein Yellow fever virus Brazil 1984 | ......................A............R...M........V........ |
| AFH35038.1:581-676 polyprotein Yellow fever virus Brazil 1987 | ......................A............R...M........V........ |
| AFH35040.1:581-676 polyprotein Yellow fever virus Brazil 1999 | ......................A............R...M........V........ |
| AFH35042.1:581-676 polyprotein Yellow fever virus Brazil 2000 | ......................A............R...M........V........ |
| AFH35044.1:581-676 polyprotein Yellow fever virus Brazil 2002 | ......................A............R...M........V........ |
| QWJ73460.1:581-676 polyprotein Yellow fever virus Brazil 2018 | ......................A............R...M........V........ |
| QWJ73461.1:581-676 polyprotein Yellow fever virus Brazil 2018 | ......................A............R...M........V........ |
| QWJ73462.1:581-676 polyprotein Yellow fever virus Brazil 2018 | ......................A............R...M........V........ |

| | |
|---|---|
| QHB50163.1:581-676 polyprotein Yellow fever virus Brazil 2018 | ......................A............R...M........V........ |
| QHB50164.1:581-676 polyprotein Yellow fever virus Brazil 2018 | ......................A............R...M........V........ |
| QHB50165.1:581-676 polyprotein Yellow fever virus Brazil 2018 | ......................A............R...M........V........ |
| QHB50166.1:581-676 polyprotein Yellow fever virus Brazil 2018 | ......................A............R...M........V........ |
| QHB50167.1:581-676 polyprotein Yellow fever virus Brazil 2018 | ......................A............R...M........V........ |
| QHB50168.1:581-676 polyprotein Yellow fever virus Brazil 2018 | ......................A............R...M........V........ |
| QHB50169.1:581-676 polyprotein Yellow fever virus Brazil 2018 | ......................A............R...M........V........ |
| QHB50170.1:581-676 polyprotein Yellow fever virus Brazil 2018 | ......................A............R...M........V........ |
| QHB50171.1:581-676 polyprotein Yellow fever virus Brazil 2018 | ......................A............R...M........V........ |
| QBK94341.1:581-676 polyprotein Yellow fever virus Brazil 2018 | ......................A............R...M........V........ |
| QBK94342.1:581-676 polyprotein Yellow fever virus Brazil 2018 | ......................A............R...M........V........ |
| UO885857.1:581-676 polyprotein Yellow fever virus Brazil 2018 | ......................A............R...M........V........ |
| QWJ73458.1:581-676 polyprotein Yellow fever virus Brazil 2018 | ......................A............R...M........V........ |
| QWJ73459.1:581-676 polyprotein Yellow fever virus Brazil 2018 | ......................A............R...M........V........ |
| AFH35034.1:581-676 polyprotein Yellow fever virus Brazil 1981 | ......................A............R...M........V........ |
| AFH35037.1:581-676 polyprotein Yellow fever virus Brazil 1984 | ......................A............R...M........V........ |
| AFH35038.1:581-676 polyprotein Yellow fever virus Brazil 1987 | ......................A............R...M........V........ |
| AFH35040.1:581-676 polyprotein Yellow fever virus Brazil 1999 | ......................A............R...M........V........ |
| AFH35042.1:581-676 polyprotein Yellow fever virus Brazil 2000 | ......................A............R...M........V........ |
| AFH35044.1:581-676 polyprotein Yellow fever virus Brazil 2002 | ......................A............R...M........V........ |
| QWJ73460.1:581-676 polyprotein Yellow fever virus Brazil 2018 | ......................A............R...M........V........ |
| QWJ73461.1:581-676 polyprotein Yellow fever virus Brazil 2018 | ......................A............R...M........V........ |
| QWJ73462.1:581-676 polyprotein Yellow fever virus Brazil 2018 | ......................A............R...M........V........ |
| QWJ73466.1:581-676 polyprotein Yellow fever virus Brazil 2018 | ......................A............R...M........V........ |
| QWJ73468.1:581-676 polyprotein Yellow fever virus Brazil 2018 | ......................A............R...M........V........ |
| QAT79253.1:573-668 polyprotein partial Yellow fever virus Brazil 2018 Aedes Albopictus | ......................A............R...M........V........ |
| UO885858.1:581-676 polyprotein Yellow fever virus Brazil 2019 | ......................A............R...M........V........ |
| QNG40777.1:581-676 polyprotein Yellow fever virus 17DD Brazil 2019 | ...I.....F.........................R..................... |

**Supp. Figure. S2:The RMSD per residue of mutant EDIII medoid structure concerning wild EDIII medoid structure of most populated cluster of 100 ns trajectories.** Thirty-four residues (296-298, 303, 313, 323, 326, 327, 331, 339-344, 353-361, 371, 380-383, 388, 392-395) show the RMSD ≥2.0 Å as compared to the wild structure. The linear red-colored dashed line represents the RMSD cut-off of 2.0 Å.

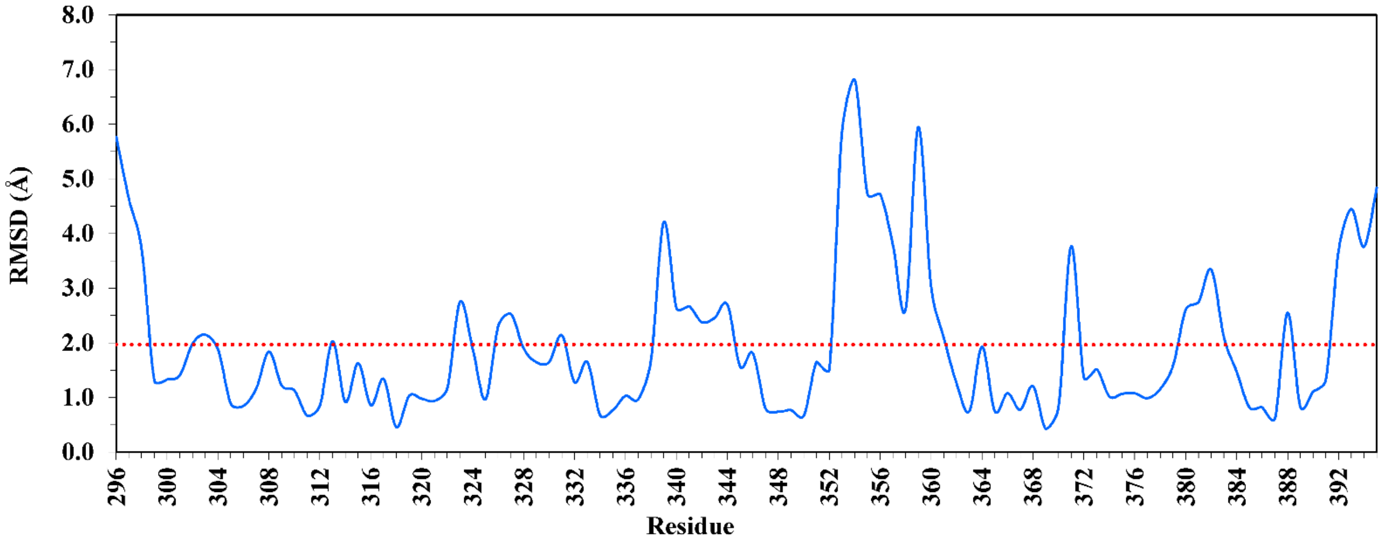


**Supp. Figure. S3:** Changes in secondary structure of each residue in wild and mutant E protein domain III (Residue 296 to 395) variants over 100 ns simulation time.  (A) Wild. (B) Mutant.

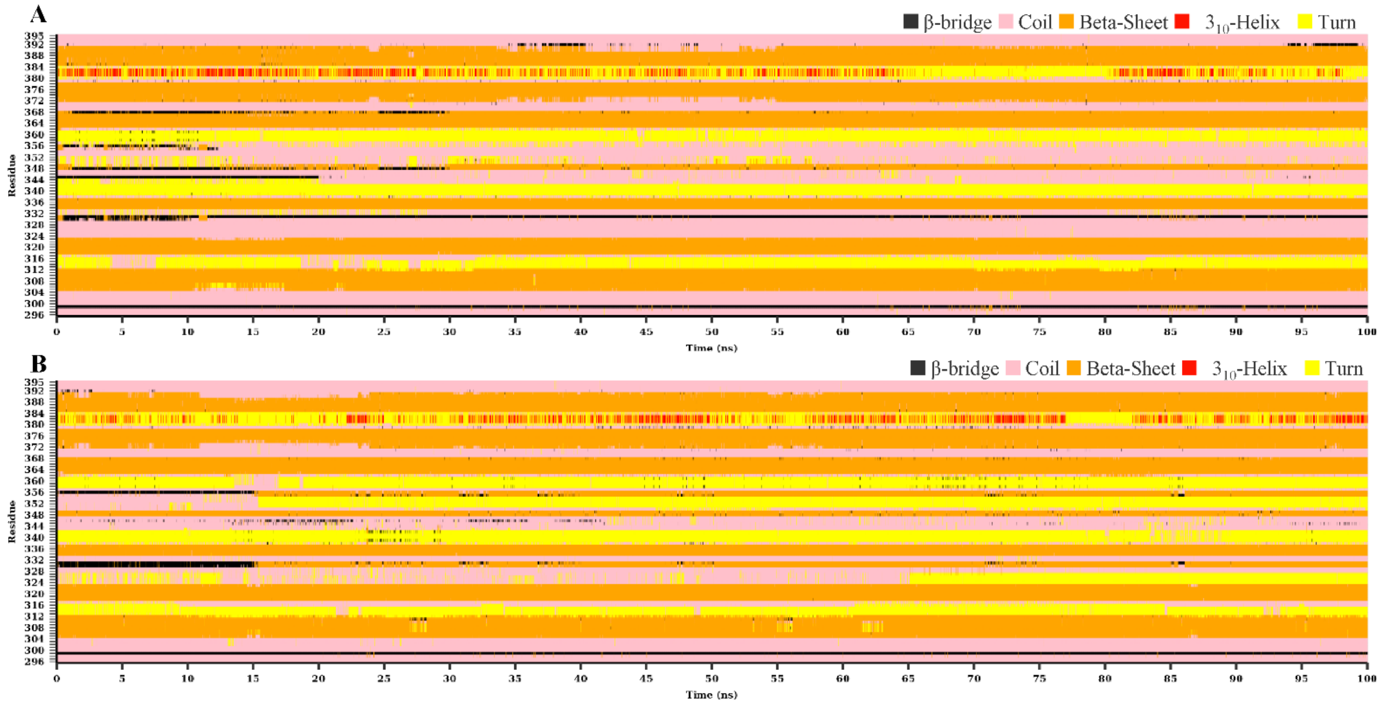

**Table S1:** The values of Cα atom's significant motion of each residues obtained from PCA across the PC1 and PC2 modes in wild and mutant EDIII trajectories.

| Residue | Wild EDII | | Mutant EDIII | |
|---|---|---|---|---|
| | PC1 | PC2 | PC1 | PC2 |
| 296 | 1.987 | 0.084 | 0.284 | 0.585 |
| 297 | 0.327 | 0.020 | 0.156 | 0.375 |
| 298 | 0.123 | 0.059 | 0.013 | 0.136 |
| 299 | 0.174 | 0.054 | 0.036 | 0.046 |
| 300 | 0.135 | 0.021 | 0.037 | 0.022 |
| 301 | 0.137 | 0.023 | 0.036 | 0.026 |
| 302 | 0.046 | 0.021 | 0.040 | 0.022 |
| 303 | 0.010 | 0.010 | 0.032 | 0.013 |
| 304 | 0.008 | 0.003 | 0.006 | 0.002 |
| 305 | 0.057 | 0.003 | 0.000 | 0.061 |
| 306 | 0.151 | 0.001 | 0.003 | 0.097 |
| 307 | 0.113 | 0.000 | 0.003 | 0.076 |
| 308 | 0.098 | 0.000 | 0.005 | 0.052 |
| 309 | 0.110 | 0.000 | 0.004 | 0.063 |
| 310 | 0.110 | 0.000 | 0.006 | 0.057 |
| 311 | 0.089 | 0.001 | 0.013 | 0.061 |
| 312 | 0.103 | 0.002 | 0.028 | 0.099 |
| 313 | 0.078 | 0.001 | 0.023 | 0.086 |
| 314 | 0.245 | 0.007 | 0.026 | 0.119 |
| 315 | 0.882 | 0.027 | 0.008 | 0.097 |
| 316 | 1.479 | 0.041 | 0.017 | 0.033 |
| 317 | 0.839 | 0.033 | 0.011 | 0.027 |
| 318 | 0.175 | 0.011 | 0.013 | 0.025 |
| 319 | 0.061 | 0.002 | 0.010 | 0.022 |
| 320 | 0.052 | 0.000 | 0.002 | 0.008 |
| 321 | 0.051 | 0.000 | 0.001 | 0.009 |
| 322 | 0.025 | 0.001 | 0.000 | 0.016 |
| 323 | 0.000 | 0.001 | 0.003 | 0.041 |
| 324 | 0.024 | 0.003 | 0.004 | 0.040 |
| 325 | 0.003 | 0.003 | 0.004 | 0.002 |
| 326 | 0.009 | 0.004 | 0.027 | 0.056 |
| 327 | 0.089 | 0.008 | 0.130 | 0.086 |
| 328 | 0.172 | 0.012 | 0.068 | 0.099 |
| 329 | 0.169 | 0.012 | 0.014 | 0.114 |
| 330 | 0.132 | 0.020 | 0.013 | 0.046 |
| 331 | 0.121 | 0.030 | 0.017 | 0.016 |
| 332 | 0.043 | 0.020 | 0.024 | 0.000 |
| 333 | 0.091 | 0.012 | 0.046 | 0.011 |
| 334 | 0.181 | 0.009 | 0.031 | 0.013 |
| 335 | 0.129 | 0.008 | 0.020 | 0.015 |
| 336 | 0.119 | 0.012 | 0.030 | 0.022 |
| 337 | 0.152 | 0.014 | 0.015 | 0.014 |
| 338 | 0.199 | 0.020 | 0.016 | 0.003 |
| 339 | 0.076 | 0.014 | 0.006 | 0.026 |
| 340 | 0.033 | 0.010 | 0.015 | 0.042 |
| 341 | 0.043 | 0.015 | 0.077 | 0.213 |
| 342 | 0.069 | 0.021 | 0.132 | 0.458 |
| 343 | 0.552 | 0.030 | 0.211 | 0.520 |
| 344 | 2.000 | 0.039 | 0.204 | 0.334 |

| | | | | |
|---|---|---|---|---|
| 345 | 1.761 | 0.034 | 0.192 | 0.216 |
| 346 | 0.303 | 0.031 | 0.083 | 0.070 |
| 347 | 0.039 | 0.027 | 0.044 | 0.015 |
| 348 | 0.077 | 0.020 | 0.028 | 0.036 |
| 349 | 0.480 | 0.015 | 0.022 | 0.038 |
| 350 | 0.964 | 0.017 | 0.008 | 0.035 |
| 351 | 1.642 | 0.011 | 0.011 | 0.050 |
| 352 | 0.994 | 0.003 | 0.062 | 0.050 |
| 353 | 0.648 | 0.007 | 0.206 | 0.071 |
| 354 | 0.538 | 0.017 | 0.320 | 0.132 |
| 355 | 0.715 | 0.024 | 0.147 | 0.134 |
| 356 | 0.762 | 0.027 | 0.036 | 0.035 |
| 357 | 0.677 | 0.029 | 0.022 | 0.082 |
| 358 | 0.475 | 0.020 | 0.011 | 0.113 |
| 359 | 0.303 | 0.015 | 0.024 | 0.064 |
| 360 | 0.305 | 0.007 | 0.025 | 0.054 |
| 361 | 0.275 | 0.005 | 0.016 | 0.116 |
| 362 | 0.177 | 0.003 | 0.003 | 0.091 |
| 363 | 0.030 | 0.001 | 0.001 | 0.025 |
| 364 | 0.005 | 0.000 | 0.000 | 0.007 |
| 365 | 0.025 | 0.000 | 0.000 | 0.004 |
| 366 | 0.070 | 0.003 | 0.000 | 0.015 |
| 367 | 0.101 | 0.010 | 0.004 | 0.019 |
| 368 | 0.091 | 0.029 | 0.013 | 0.026 |
| 369 | 0.112 | 0.048 | 0.027 | 0.015 |
| 370 | 0.328 | 0.035 | 0.050 | 0.015 |
| 371 | 0.266 | 0.018 | 0.063 | 0.011 |
| 372 | 0.135 | 0.005 | 0.092 | 0.006 |
| 373 | 0.080 | 0.015 | 0.073 | 0.003 |
| 374 | 0.080 | 0.017 | 0.031 | 0.003 |
| 375 | 0.077 | 0.008 | 0.013 | 0.003 |
| 376 | 0.103 | 0.005 | 0.005 | 0.005 |
| 377 | 0.116 | 0.005 | 0.001 | 0.007 |
| 378 | 0.235 | 0.007 | 0.006 | 0.012 |
| 379 | 0.365 | 0.014 | 0.032 | 0.011 |
| 380 | 0.401 | 0.022 | 0.051 | 0.019 |
| 381 | 0.425 | 0.030 | 0.038 | 0.019 |
| 382 | 0.283 | 0.064 | 0.005 | 0.009 |
| 383 | 0.184 | 0.114 | 0.042 | 0.009 |
| 384 | 0.224 | 0.063 | 0.026 | 0.016 |
| 385 | 0.218 | 0.014 | 0.016 | 0.017 |
| 386 | 0.182 | 0.007 | 0.010 | 0.018 |
| 387 | 0.182 | 0.004 | 0.002 | 0.014 |
| 388 | 0.165 | 0.007 | 0.012 | 0.016 |
| 389 | 0.110 | 0.014 | 0.024 | 0.012 |
| 390 | 0.112 | 0.029 | 0.010 | 0.021 |
| 391 | 0.207 | 0.063 | 0.021 | 0.087 |
| 392 | 0.502 | 0.066 | 0.046 | 0.333 |
| 393 | 0.840 | 0.287 | 0.181 | 0.793 |
| 394 | 1.003 | 1.017 | 0.543 | 1.308 |
| 395 | 1.566 | 2.000 | 2.000 | 2.000 |
